## Supplementary figures and images for "Human disturbance increases spatiotemporal associations among mountain forest terrestrial mammal species"

### Figure 3-figure supplement 1

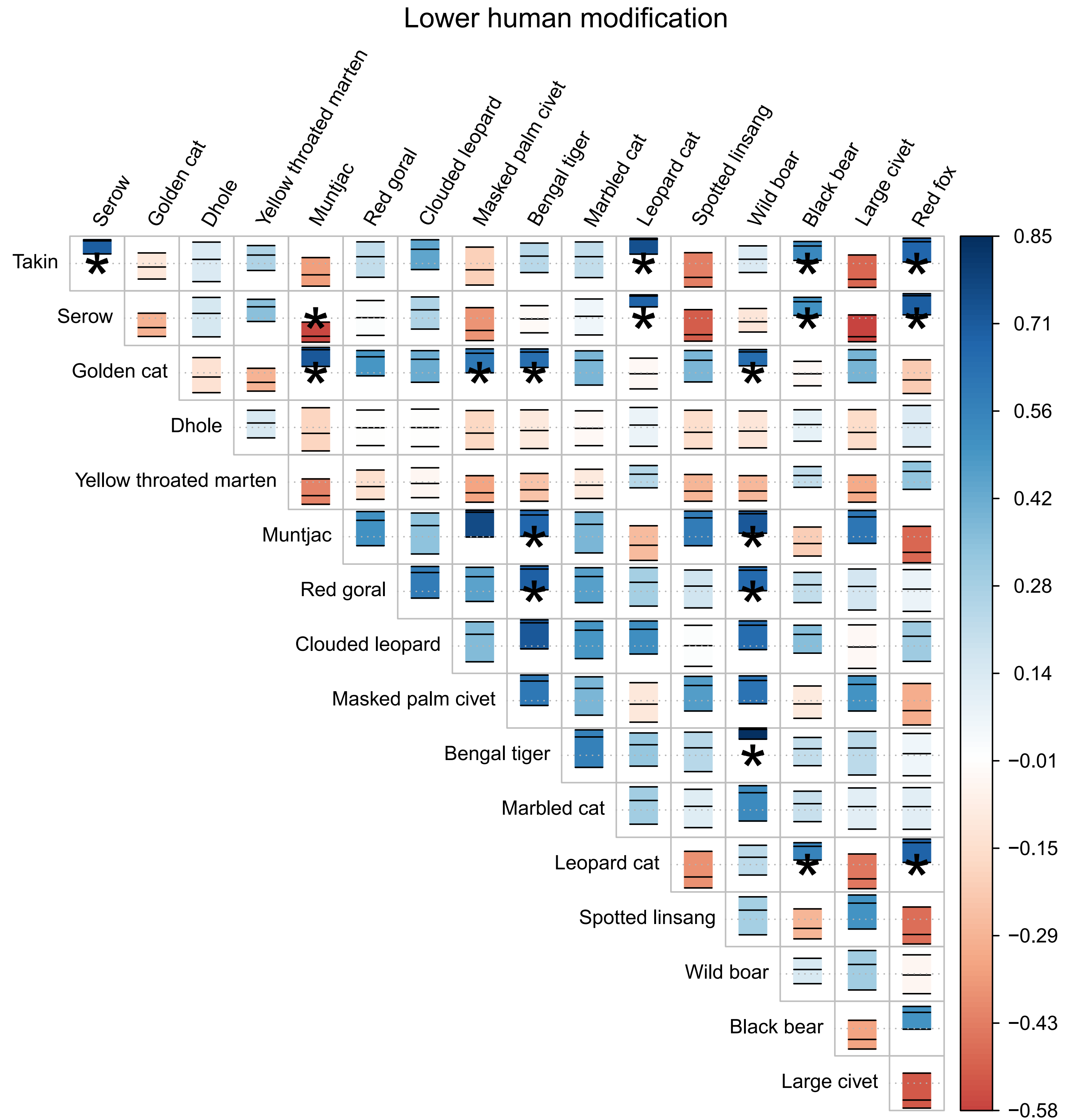

### Figure 3-figure supplement 2

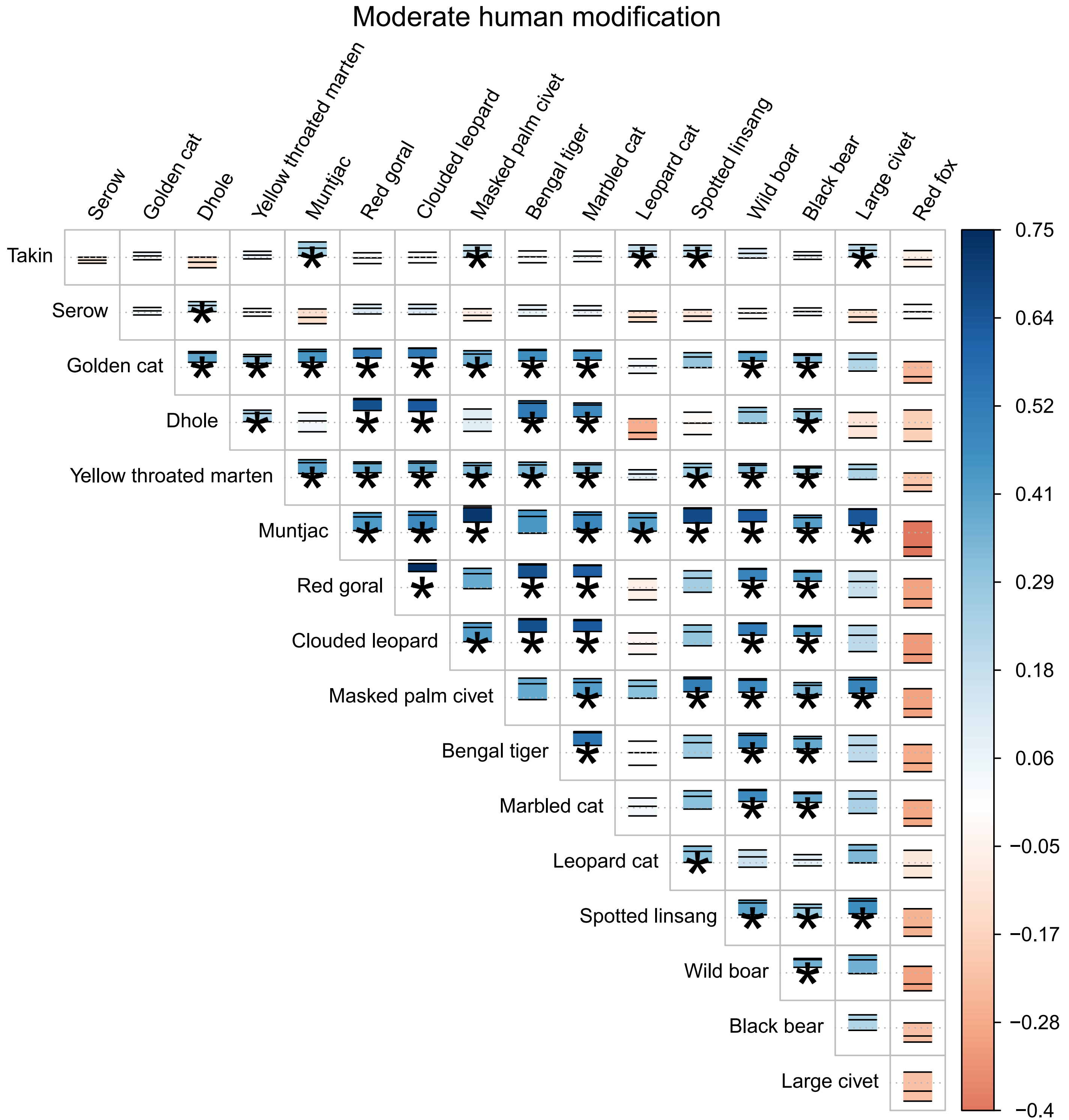

### Figure 3-figure supplement 3

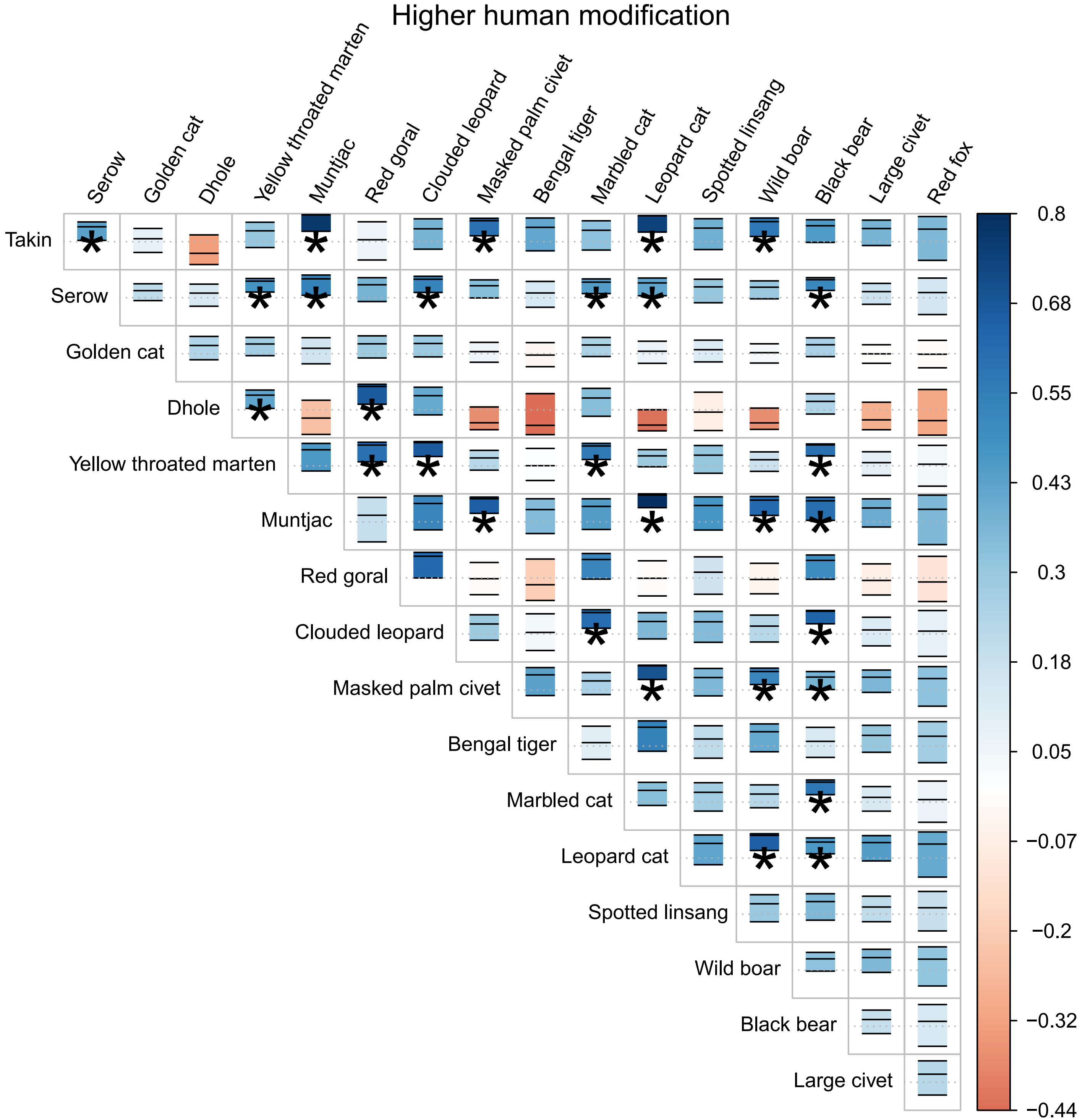

### Figure 4-figure supplement 1

#### Lower human presence

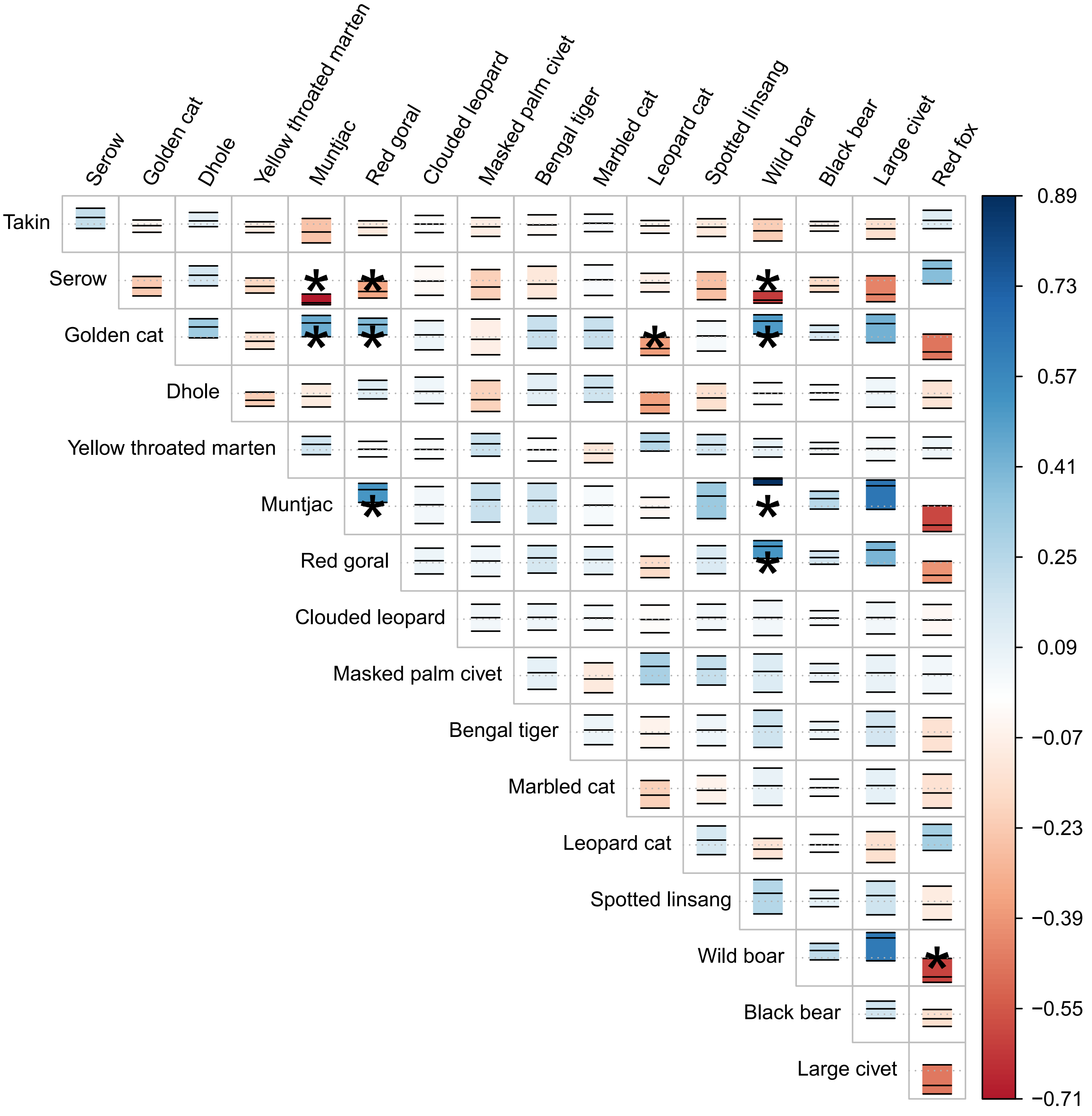

### Figure 4-figure supplement 2

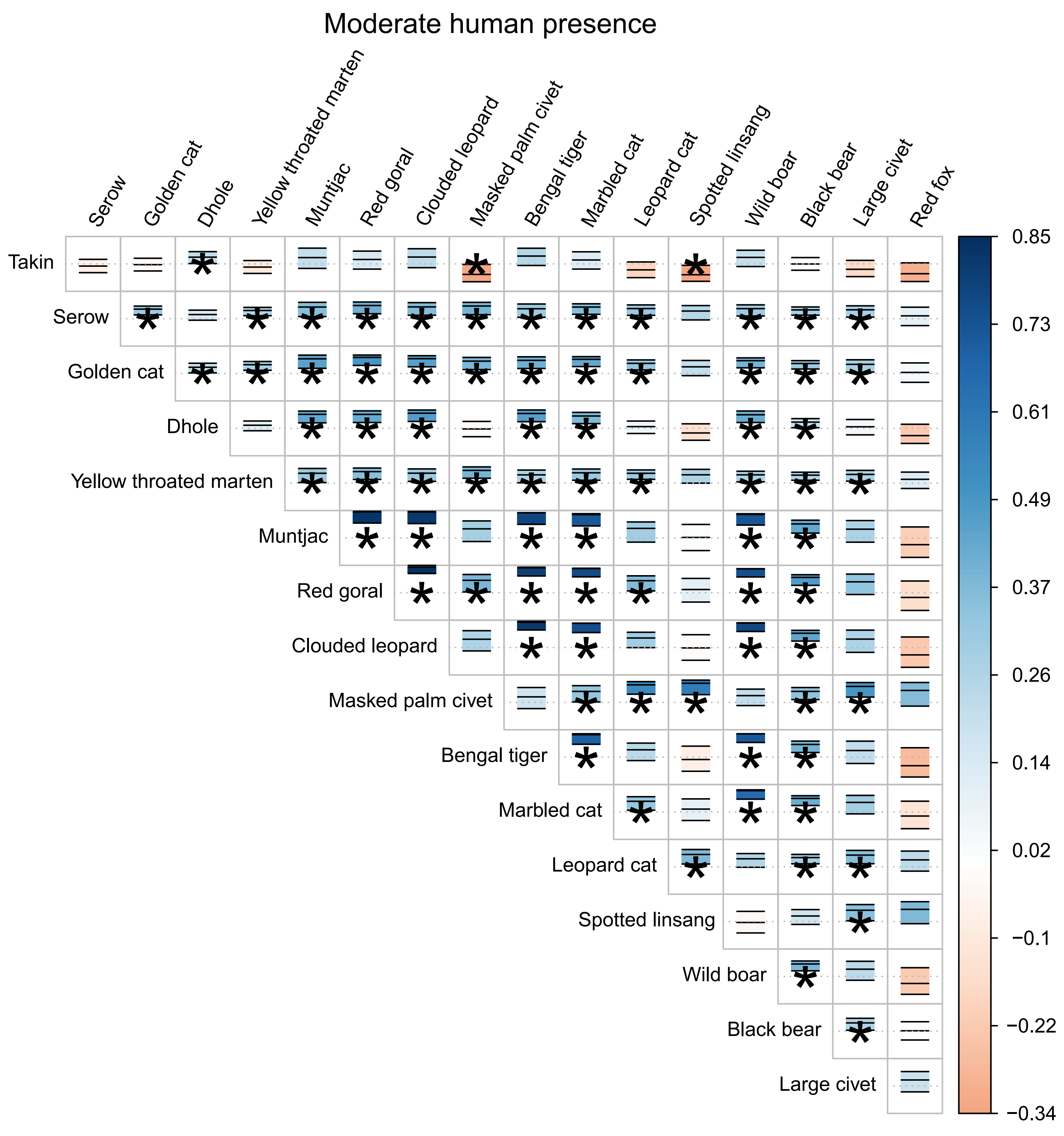

### Figure 4-figure supplement 3

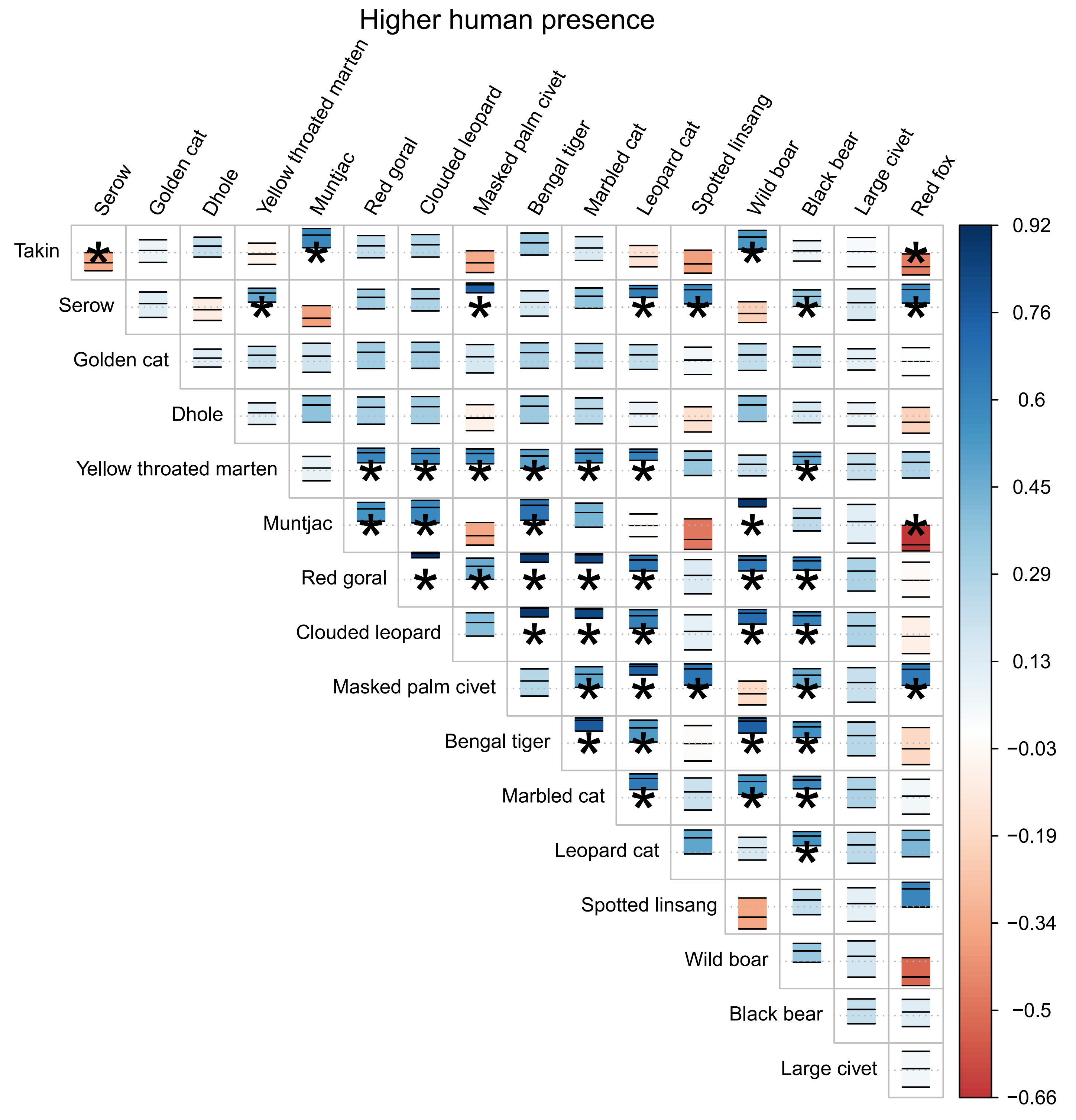
